# Damage-induced phosphorylation of BRC-1/BRD-1 in meiosis preserves germline integrity

**DOI:** 10.1101/2025.03.30.646171

**Authors:** N Fernández-Fernández, M Chacón, LP Camino, T García-Muse

## Abstract

Multiple DNA repair pathways have evolved to safeguard genome integrity and ensure organismal viability in the face of DNA damage. Errors in DNA repair processes in meiosis can lead to aneuploidy and developmental defects, but the processes that protect the germline from DNA damage remain poorly understood. Here we report a DNA damage-induced phosphorylation of the BRC-1/BRD-1 heterodimer that is essential for germline integrity in *C. elegans*. Failure to phosphorylate BRC-1/BRD-1 in response to DNA damage results in meiotic DSBs accumulation, chromosome breakage, catastrophic diakinesis and loss of fecundity. We further show that these defects are driven by the activity of *C. elegans* Bloom and Mus81, which catalyse Holliday junction dissolution and resolution, respectively. Hence, we propose that phosphorylation of BRC-1/BRD-1 in response to excess meiotic DSBs constitutes a key regulatory step that ensures the proper resolution of recombination intermediates required to preserve germline integrity.

## Introduction

The tumour suppressor proteins BRCA1 (Breast Cancer 1) and BARD1 (BRCA1-associated RING domain protein 1) perform roles in replication fork protection, checkpoint signalling and DNA repair by homologous recombination (1). Mutations in *BRCA1* and *BRCA2* have been linked to an increased lifetime risk of developing certain types of cancer, including breast, ovarian and prostate cancers in humans (2, 3). A body of evidence has implicated BRCA1 in regulating the resection of DNA double-strand breaks (DSBs), whereas BRCA2 facilitates the recruitment and initial nucleation of the Rad51 recombinase onto processed DSBs. During meiosis, programmed DSBs are generated to initiate homolog recombination (HR), which is required to promote the formation of inter-homolog crossovers (COs) needed for faithful chromosome segregation. These joint DNA molecules must disengage in order to segregate, and this is achieved by redundant endonucleases (called resolvases) and the BTR complex (4). In *C. elegans* MUS-81 and SLX-1 have overlapping roles with the Bloom syndrome helicase ortholog, HIM-6 in processing recombination intermediates (5–7). The *C. elegans* Bloom ortholog HIM-6 suppresses heterologous recombination in the germline, which could lead to translocations and genome rearrangements (8).

Studies in mice have shown that hypomorphic mutations or deficiencies in BRCA1 lead to defects in meiotic recombination, resulting in chromosome abnormalities and impaired fertility (9). In *C. elegans* BRC-1 and BRD-1 have also been implicated in meiosis and DNA repair during meiotic recombination. The BRC-1/BRD-1 complex localizes to the synaptonemal complex (SC) in an interdependent manner (10, 11). BRC-1 and BRD-1 are dispensable for meiotic crossover formation but are required for meiotic homolog-independent DSB repair (12, 13). Under defective meiosis BRC-1 and BRD-1 mediates RAD-51 filament stability and crossover formation (10–12). Moreover, it has been shown that BRC-1 and BRD-1 prevent recombination between heterologous templates and repress error-prone DSB repair through non-homologous end joining (NHEJ) and polymerase theta-mediated end joining (TMEJ) (14–16). The mechanisms that regulate BRC-1 and BRD-1 activation after exogenous DNA damage in meiosis remain unclear.

The DNA-damage activated kinases ATM (ataxia-telangiectasia-mutated) and ATR (ataxia-telangiectasia-related) play central roles in DSB sensing and repair in mitotic cells (17–19). ATM and ATR kinases also localize to meiotic chromosomes and have been implicated in promoting homologous recombination (HR), repair-template choice and crossover (CO) control (20). In *C. elegans* the ATR kinase, ATL-1, is essential for mitotic cell cycle arrest and induction of apoptosis in response to DNA damage but shows no obvious meiotic defects in SC assembly (21). Conversely, *C. elegans* ATM, ATM-1 is necessary for the restoration of chromatin and re- synapsis of the chromosome axes after irradiation (22). ATM-1 seems to bias the choice of repair template to the homologous chromosome, with both checkpoint kinases acting redundantly to promote RAD-51 accumulation at DSBs (23). Recently, both kinases have been shown to function at meiotic entry to reshape the genome through cohesins to promote interhomolog interactions and meiotic recombination (24). Specifically, ATM and ATR phosphorylate components of the SC to maintain architectural integrity, promoting DSB repair pathway choice to avoid NHEJ (25, 26).

Here we investigated how exogenous DNA damage is repaired in the meiotic germline of *C. elegans*. We present evidence that excessive DSBs result in IR-dependent phosphorylation of the BRC-1 and BRD-1 heterodimer, which regulates the correct resolution of recombination intermediates. We identify a cluster of S/T-Q motifs that form DNA damage-induced phosphorylation sites in the BRC-1 and BRD-1 proteins. The corresponding non-phosphorylatable (BRC-1^4A^ and BRD-1^3A^) mutants exhibit IR sensitivity and defects in DSB repair that it is exacerbated by exogenous DNA damage, demonstrating that failure to phosphorylate the heterodimer BRC-1/BRD-1 impairs proper DSB repair. Surprisingly, we show that phosphorylation is dispensable for inter-homolog DSB repair but is required to avoid inappropriate processing of recombination intermediates by Bloom and Mus81. Hence, our data reveal a mechanism by which the meiotic checkpoint acts to protect the germline from exogenous DNA damage and genetic instability.

## Materials and methods

### Experimental model maintenance and *in vivo* assays

#### Strains and maintenance

Standard methods were used for the maintenance and manipulation of C. elegans strains (27, 28). Nematode strains were provided by the Caenorhabditis Genetics Center, which is funded by the NIH National Center for Research Resources, except for 1960 ollas strain, *polq-1, mus-81* and *him-6* knock-out strains, generated and/or kindly provided by Verena Jantsh group (11). The strains with transgenic *brc-1^(4A)^*, *brd-1^(3A)^*and *polq-1;brc-1^(4A)^* alleles were generated using CRISPR/Cas9 (29, 30) by microinjection into N2(wt). The double mutants *brc-1^(4A)^;GFP::msh-5, brd-1^(3A)^;GFP::msh-5, mus-81;brc-1^(4A)^, him-6;brc-1^(4A)^, msh-5;brc-1^(4A)^*and *syp-1;brc-1^(4A)^*, were generated by crossing the corresponding strains. All strains are listed in Table S2.

#### Embryonic lethality

Embryonic lethality was scored by comparing the number of eggs that hatch to produce viable progeny versus the total number of eggs laid. Briefly L4 hermaphrodites grown at 20°C were individually plated. The animals were transferred to new plates once every 24 hours until the egg laying stopped. Eggs laid were immediately counted. When each brood reached adulthood, the total number of live animals per brood was counted and checked against the egg count to give the total brood size and an estimate of the embryonic lethality frequency. The number of larval arrested and male progeny animals was also noted. A minimum of three experiments were performed for each strain. The total number of single hermaphrodites analyzed is indicated in Table 1.

**Table 1.** Viability Analysis of *brc-1^(4A)^* and *brd-1^(3A)^* Mutant Alleles.

| Genotype | Average Brood + SD (n)<sup>a</sup> | Percentage Viable Embryos (n)<sup>b</sup> | Percentage Male (n)<sup>c</sup> |
| --- | --- | --- | --- |
| N2 (WT) | 292.73 + 16.05 (19) | 98.69 (5649) | 0.04 (5598) |
| *brc-1*<sup>(4A)</sup> | 297.33 + 12.39 (11) | 100 (3262) | 0.03 (3262) |
| *brd-1*<sup>(3A)</sup> | 290.5 + 25.27 (8) | 99.37 (2353) | 0 (2338) |

a. Parentheses indicate the total number of singled hermaphrodites for wich entire brood size were scored.
b. Parentheses indicate the total number of fertilized eggs scored.
c. Parentheses indicate the total number of adults scored.

For brood analysis after irradiation, L4 animals were exposed to the indicated Gy doses of γ-ray from BioBeam8000. After 24 or 48 hours, five post-irradiation P0 worms were plated to lay eggs for 10 hours. 36 hours later the number of hatched F1 larvae, dead embryos and males were counted (31). Three plates were counted for each strain and condition, and the experiment was repeated four times.

### Generation of nematode strains

#### Generation of brd-1 non-phosphorylable mutant and polq-1;brc-1^(4A)^ double mutant by <u>CRISPR/Cas9 genome editing</u>

The generation of *brc-1^(4A)^* mutant was ordered to SunyBiotech. *brd-1^(3A)^* mutant worms were generated in a 2 steps genome edition by CRISPR/Cas9 as in (30). N2 was used for injection and *dpy-10* co-edition was used as a positive control of Cas9 activity. Injection mixes contained Cas9 (250 ng/ml), ALT-R tracrRNA (141.97 ng/ml), ALT-R *dpy-10* crRNA (14.39 ng/ml), ALT-R *target gene* crRNA (59 ng/ml), ssODN *dpy-10(cn64)* repair template (28 ng/ml), ssODN *target gene repair template* (116 ng/ml) and nuclease-free H_2_O. *polq-1;brc-1^(4A)^* mutant worms were generated also by CRISPR/Cas9. In this case, *brc-1^(4A)^* was used for injection and two different ALT-R *target gene* crRNA were used in the injection mix. General reagents were acquired from IDT and are listed in Table S3. The resulting transformants (roller or dumpy phenotype worms) were transferred to new plates and genotyped. All primers and RT are listed in Table S4.

#### Worm genotyping

The resulting transformants were checked by single worm PCR using MyTaq DNA-polymerase and restriction enzymes digestion. gDNA was obtained by single worm lysis and used in PCRs. Restriction enzymes digestion was used to check for the integration of the *brd-1^(3A)^* repair template (Biolabs restriction enzymes PstI and PvuII are listed in Table S3). Homozygote phospho-alleles candidates were sequenced to check the integration of the expected mutations and resultant strains were back crossed twice.

The double mutant strains were also checked by single worm PCR (primers are listed in Table S4). To check for *brc-1^(4A)^* and *brd-1^(3A)^* alleles in gfp::*msh-5* homozygote candidates, a fragment of the respective gene was amplified by PCR and sequenced. The presence of gfp::*msh-5* allele was determined by taking advantage of the fluorescence the mutation generates. To check for *brc-1^(4A)^* allele in *polq-1;brc-1^(4A)^, mus-81;brc-1^(4A)^, him-6;brc-1^(4A)^, msh-5;brc-1^(4A)^*and *syp-1;brc- 1^(4A)^*, specific primers for *brc-1^(4A)^*and brc-1 wild-type alleles amplification were designed. Restriction enzymes digestion was used to check for the presence of *msh-5* and *syp-1* knock-out alleles (Biolabs restriction enzymes Hyp188I and BstAPI are listed in Table S3). The primers used for all strains genotyping are listed in Table S4.

### Molecular Biology

#### sRNA guides design

To design the sgRNA recognition sites and the repair templates for *brd-1^(3A)^* mutant, we followed (29) protocol. To select PAM sites, we used CRISPOR website (http://crispor.tefor.net/). The designed sRNA guides are indicated in Table S4.

#### Peptide arrays and kinase assays

For the peptide array studies, 18-mer peptides were made by solid-phase synthesis and purified by high-performance liquid chromatography, and their sequences were verified by mass spectroscopy. The 18-mer peptides peptides juxtaposed by three amino acids until scanning the complete BRC-1 and BRD-1 proteins. All peptides contained an N-terminal biotin group with an aminohexanoic spacer to be spotted onto cellulose membrane. The membrane was activated by soaking in methanol for 2 min and washed twice with kinase buffer supplemented with 3% BSA. *In vitro* phosphorylation was performed by incubating the membrane in 5 mL of kinase buffer supplemented with N2 worm extracts (protein concentration of 10 mg/ mL) and 100 μCi of [32P] γ-ATP. After adding stop buffer, the membrane was washed sequentially in 1 M NaCl, then 1% SDS, and finally 0.5% phosphoric acid solution. After washing in 96% ethanol, the membrane was dried and exposed to autoradiography film.

#### Gene knockdown

For gene knockdown assays, RNAi depletion by the feeding method was performed as described in (32), with modifications. HT115 bacteria containing the pL4400 vector or the correspondent RNAi feeding construct were seeded on LB plates containing ampicillin and tetracycline and incubated overnight at 37 °C. A bacteria colony was suspended in 2 ml of LB, containing ampicillin, and incubated overnight at 37 °C. The bacteria inoculum was seeded in NGM 6-wells plates, with ampicillin, tetracycline and 6mM isopropylthio-b-D-galactosidase and incubated at RT for 36 hours. 10 late L4 worms for each strain and condition were transferred to wells, and irradiated at 75Gy. Worms were incubated for 48h at 20 °C before DAPI staining of the germlines.

### Cellular biology

#### Immunostaining

For RAD-51 and SYP-1 immunostaining worms were treated as described in (33), with modifications. One day post-L4 adult gonads were dissected in TBSTw 0.1% (1 TBS, 0.1% Tween20) on a Superfrost Plus slide (VWR), and were fixed for 5 minutes in 1% paraformaldehyde. Gonads were then flash frozen in liquid N_2_ and the cover slip was removed. The slide was placed in -20°C MeOH for 10 min and washed 3 times in TBSTw 0.1% for 10 min before being placed in block (1 TBS, 0.1% Tween20, 0.7% Bovine Serum Albumin) for 1 hour. Slides were incubated overnight at 4°C in a dark humidifying chamber with diluted primary antibody to stain. Dilutions used were rabbit α-RAD-51 (1:5000) and guinea pig α-SYP-1 (1:500) in TBSTw 0.1%. Next day gonads were rinse and then washed 3 times in TBSTw 0.1%, each for 10 minutes at RT, and incubated for 2 hours with the secondary antibody in TBSTw 0.1% (α-RABBIT 1:200, α-GUINEA PIG 1:500) in a dark humidifying chamber. Gonads were rinse and 1µg/ml DAPI in TBSTw 0.1% was added to each slide and incubated for 10 min. Slides were washed 4 times in TBSTw 0.1% for 10 min and mounted with Vectashield. They were maintained at 4°C for 2-3 days prior to imaging. For irradiation experiments, one day post-L4 adults were irradiated with 75Gy dose of γ-ray from BioBeam8000 and 48 hours post-irradiation gonads were dissected for immunostaining as described.

For BRC-1 and BRD-1 antibody worms were treated as described in (25), with modifications. 4 hours after irradiation worms were dissected in PBS on poly-lysine slides, fixed for 15 minutes in 4% paraformaldehyde and replaced for 5 minutes in TBSBTx (TBSB +0.1% TX100). The slides were washed twice for 10 minutes with TBSTw 0.1% and one more for 30 minutes with TBSB (TBS + 0.5% BSA). They were incubated overnight at 4 °C with a BRC-1 or BRD-1 (1:100) and SYP-1 (1:300) antibody dilution. Next day gonads were rinse and then washed 3 times in TBSTw 0.1%, each for 10 minutes at RT, and incubated for 2 hours with the secondary antibodies (α- RABBIT 1:500 and α-CHICKEN 1:500). Gonads were rinse and then washed three times for 10 minutes with 2mg/ml DAPI in TBSTw 0.1% and mounted with 10 mL Vectashield (with 1 mg/ml DAPI) per sample for further analysis.

#### SYTO12 for apoptotic corpses quantification

For apoptotic corpses (AP) analysis, 24 hours post-L4 animals were incubated in the dark with a 40mM aqueous solution of the genotoxic agent SYTO-12. After 4 hours, to facilitate the apoptotic corpses visualization, worms were transferred to new dishes with OP50 bacteria and were incubated for 1 hour at 25°C, to metabolize the excess of SYTO-12 present in the intestine. After the indicated time worms were transferred to slides with agarose pad to score under the microscope the presence of fluorescent bodies indicating cells undergoing apoptosis (31). The experiment was repeated a minimum of three times and a minimum of 30 total worms for each strain were scored. For irradiation experiments, L4 worms were irradiated with 75Gy dose of γ-ray from BioBeam8000 and 12-, 24- and 36-hours post-irradiation gonads were treated as described above.

#### GFP::MSH-5 foci quantification

One day post-L4 adults were irradiated with 75Gy dose of γ-ray from BioBeam8000. 48 hours post-irradiation, GFP::MSH-5 foci were detected as described in (34), with modifications. For GFP::MSH-5 detection, worms were dissected in TBSTw 0.1% and directly frozen in liquid nitrogen. After freeze-cracking, slides were incubated in methanol at −20°C for 5 min. Gonads were immediately washed with TBSTw 0.1% for 5 min and fixed with 4% PFA in 100 mM K_2_HPO_4_. Slides were washed three times in TBSTw 0.1% for 10 min, and incubated with g/ml DAPI in TBSTw 0.1% for 10 min. Slides were mounted with Vectashield and germlines were examined with fluorescence microscopy. 10-12 nuclei were counted for late-pachytene region for a minimum of four germlines per genotype and condition.

#### Diakinesis nuclei quantification

For visualization and quantification of diakinesis nuclei, germlines were dissected and stained with DAPI as described for GFP::MSH-5 detection, except that paraformaldehyde was used at 1% and fixation was performed before liquid nitrogen freeze-cracking. DAPI bodies from a minimum of 2- 4 diakinesis nuclei were counted or morphologically checked for 10-15 germlines per genotype and condition.

#### Fluorescence microscopy

Leica DM6000B was used to examine the germlines with 100X HCXPL-APO/1.40 OIL lens, and images were captured using Leica LAS-AF computer software for RAD-51 and SYP-1 immunostaining and GFP::MSH-5 foci quantification. Between 150-220 confocal planes of 0.2 μm- distance were taken per each gonad (depending on the thickness of the gonad) and non-saturated laser conditions were adjusted for each experiment. Analysis was performed for half of the germline along the dorsal-ventral axis, using the maximum projection for RAD-51 quantification. Nikon SMZ-645 was used to examine the germlines with 60X PL-APO/1.45 OIL lens for apoptotic corpse analysis.

For BRC-1 and BRD-1 immunostaining and diakinesis nuclei quantification Zeiss AxioImager.2 was used to examine the germlines with Plan-Apochromat 63x and 100x /1.40 oil lens, and images were captured using Zeiss Zen2 computer software. Between 70-130 confocal planes of 0.2 μm- distance were taken per each gonad (depending on the thickness of half the gonad or diakinesis stage) and non-saturated laser conditions were adjusted for each experiment.

#### RAD-51 foci quantification

Analysis of RAD-51 foci was performed as described in (33). Each germline was divided into 6 regions, corresponding to mitosis division zone, transition zone, early pachytene, medium pachytene and late pachytene zones, and diplotene zone. The number of foci per nucleus in each region of the germline were quantified. A minimum of four germlines per genotype and condition were analyzed. Data shows the % of nuclei of the different categories based in the number of foci/nuclei. A minimum of four germlines per genotype were analyzed.

#### Quantification and statistical analysis

Quantification was performed always in raw images. After quantification, for beauty purposes in the images shown a background subtraction plugging was applied using FiJI software. Statistical significance was determined with unpaired t-test 1way ANOVA or 2way ANOVA using PRISM software (Graphpad Software Inc.). Specific replicate numbers (n) for each experiment can be found in the corresponding figure legends. In all figures, means are plotted and standard deviation (SD) or standard error of the mean (SEM) is represented as error bars.

## Results

### DNA damage phosphorylation sites in BRC-1- and BRD-1

We have previously described a DNA damage-induced phosphorylation of the SC component SYP-1 that channels the repair of excessive DSBs through the activity of BRC-1 (25). Given the importance of BRCA1 in regulating pathway choice in mitotic cells, we considered the possibility that phosphorylation of BRC-1 and/or BRD-1 may play an analogous regulatory role during meiotic HR repair. To explore this possibility, we examined damage-induced phosphorylation events on BRC-1 and BRD-1 using peptide arrays against both proteins. Protein extracts from untreated worms or worms treated with 75Gy of IR were used for *in vitro* kinases assays, which revealed DNA damage-induced phosphorylation events on both BRC-1 and BRD-1. Specifically, we identified a SQ motif surrounded by several serine and threonine residues between amino acids 436 and 443 within the BRCT domain of BRC-1 (Figure 1A). In the case of BRD-1, we identified a cluster of three TQ motifs within 104-132 amino acids adjacent to the RING domain (Figure 1B). Both BRCT and RING domains of the worm proteins are highly conserved with human and xenopus orthologs BRCA1 and BARD1 (35).

**Figure 1.**
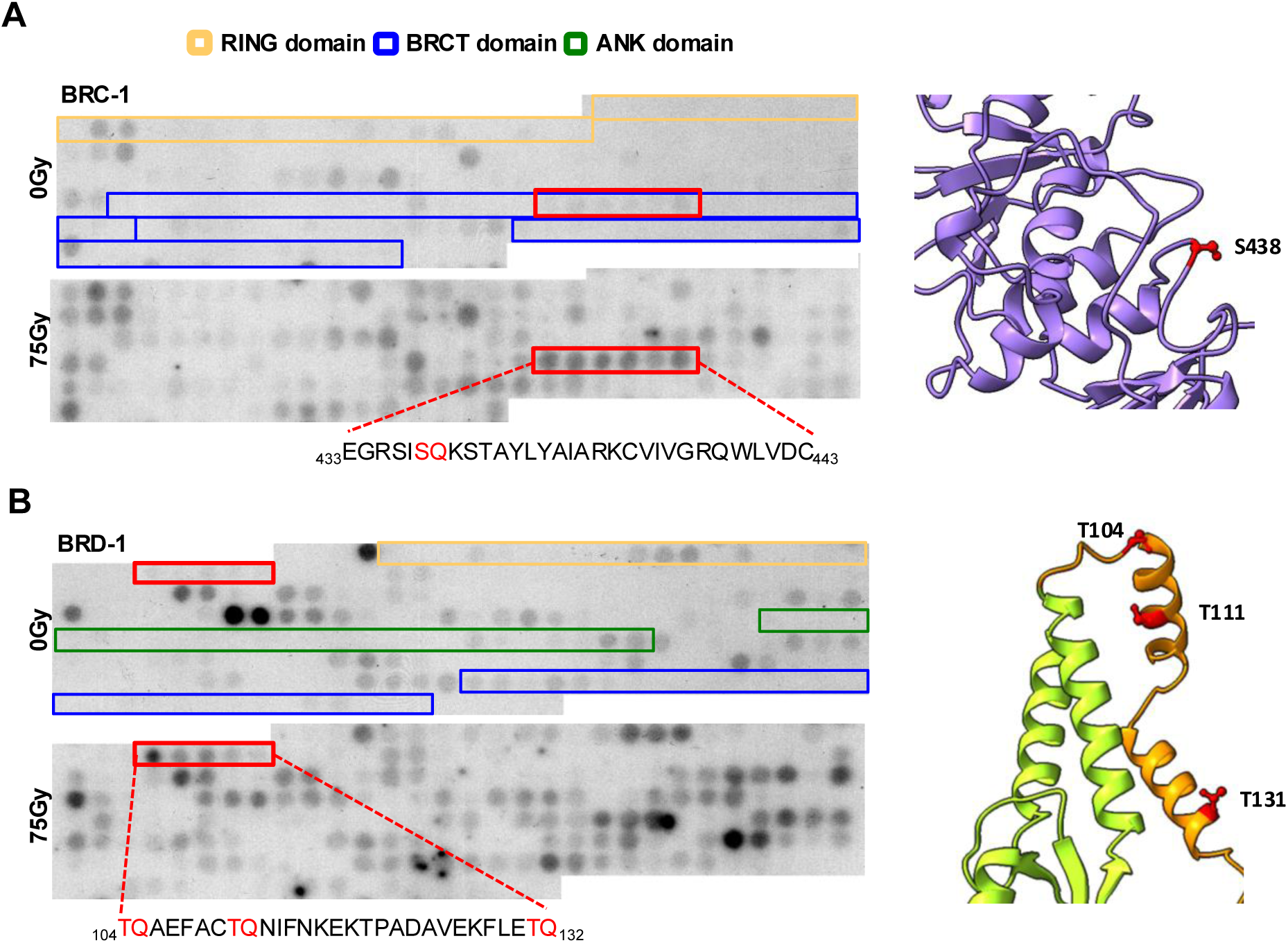
*In vitro* DNA damage phosphorylation of BRC-1 and BRD-1 proteins. **(A)** Left, the BRC-1 peptide array with N2(WT) protein extracts without DNA damage (top) and with N2(WT) extracts after 75 Gy (bottom). Each of the spots represents an 18-mer peptide fragment juxtaposed by three amino acids (aa) scanning the complete BRC-1 protein. Positive serial spots (detected by autoradiography) corresponding to the specific DNA damage-phosphorylated region are boxed in red. Scheme depicts the phosphorylation site established by the peptide array data, with the possible phosphorylation residues highlighted in red. Right, the predicted structure of BRC-1 BRCT domain determined by AlphaFold. In red, the putative phosphorylated S/T-Q motif for BRC-1 (S438). **(B)** Left, the BRD-1 peptide array with N2(WT) protein extracts without DNA damage (top) and with N2(WT) extracts after 75 Gy (bottom). Each of the spots represents an 18-mer peptide fragment juxtaposed by three amino acids (aa) scanning the complete BRD-1 protein. Positive serial spots (detected by autoradiography) corresponding to the specific DNA damage-phosphorylated region are boxed in red. Scheme depicts the phosphorylation site established by the peptide array data, with the possible phosphorylation residues highlighted in red. Right, the AlphaFold predicted structure of BRD-showing the RING domain (green). In red, the putative phosphorylated residues for BRD-1 (T104, T111 and T131).

To examine the structural context of the phospho-motifs we used ChimeraX to predict the structure of *C. elegans* BRC-1 and BRD-1 BRCT and RING domain, respectively (36). We used the predicted structures determined by AlphaFold of BRC-1 BRCT domain (Figure 1A), and BRD- 1, focusing on the RING domain (green) and its continuous region (orange), where the motifs of interest are located (Figure 1B). The predicted conformations show that all phospho-motifs are solvent exposed on the surface of the predicted protein structure.

To investigate the biological relevance of the damage-induced phosphorylation sites in BRC-1 and BRD-1 proteins, we used CRISPR-Cas9 to generate non-phosphorylatable alleles in which the putative phosphorylated residues were changed to alanine. For BRC-1 we generated the S438A mutation in addition to S436A, S441A and T442A mutations to ensure there was no possibilities for compensatory phosphorylation. This resulted in the *brc*-*1*^4A^ allele. For BRD-1 we generated the *brd*-*1*^3A^ in which we changed each of the threonines (T104, T111 and T131) to alanine. To eliminate possible off-targets of Cas9, the resulting transgenic lines were then back- crossed with the N2(wt) and *brd-1* knockout, respectively. We first analysed the effect of these mutations on worm development by performing viability assays. Similar to what has been described for the loss of *brc-1* or *brd-1* (12, 35), both mutated alleles are viable and do not show overt differences in brood size with respect to the N2 (wt) (Table 1). This data indicates that disruption of BRC-1 and BRD-1 phosphorylation does not negative impact development.

### The lack of BRC-1 and BRD-1 phosphorylation results in mild meiotic DSB repair defects

In previous studies, it was shown that the loss of *brc-1* or *brd-1* leads to a mild meiotic defect, supported amongst other observations by the appearance of males, which results from loss of an X-chromosome (12, 35). During the viability assays we did not observe an increase of males in the progeny of the *brc*-*1*^4A^ and *brd*-*1*^3A^ mutants (Table 1). Nevertheless, we also performed a cellular analysis of the germline, which allows for temporal and spatial analyses of meiotic progression through prophase I (37), with markers of the key steps in meiosis. Homologous chromosome synapsis can be studied by immunostaining of the SC central region protein SYP-1 (38); using an antibody against SYP-1 we observed that its localization between paired homologous chromosomes in the *brc*-*1*^4A^ and *brd*-*1*^3A^ mutant strains is indistinguishable from the N2(wt) (Supplementary Figure S1), indicating that synapsis is unaffected in these mutants.

Next, we studied programmed meiotic DSBs repair using an antibody against RAD-51, which catalyses the strand invasion and exchange steps during HR (39). During normal meiosis in N2(wt) worms, RAD-51 foci are observed at sites of SPO-11-induced meiotic DSBs, with RAD- 51 foci first appearing in the transition zone and progressively increasing to a maximum number in mid-pachytene and finally disappearing in late pachytene (Figure 2A, 2B) (38). In the case of *brc*-*1*^4A^ and *brd*-*1*^3A^ mutant germlines, we observed that RAD-51 foci start to appear in the transition zone but are slightly increased when compared to N2(wt) germlines (Figure 2A, 2B). This was more prominent in *brc*-*1*^4A^ where at later regions we observed 27% of nuclei with persistent RAD-51 foci at unrepaired DSBs. Since the accumulation of unrepaired DNA damage leads to apoptosis, we also scored germ cell apoptosis and found that apoptotic corpses were significantly increased in *brc*-*1*^4A^ and *brd*-*1*^3A^ mutant strains when compared to N2(wt) (Figure 2C). This data suggests that the inability to phosphorylate BRC-1 or BRD-1 leads to mild defects in the repair of programmed meiotic DSBs.

**Figure 2.**
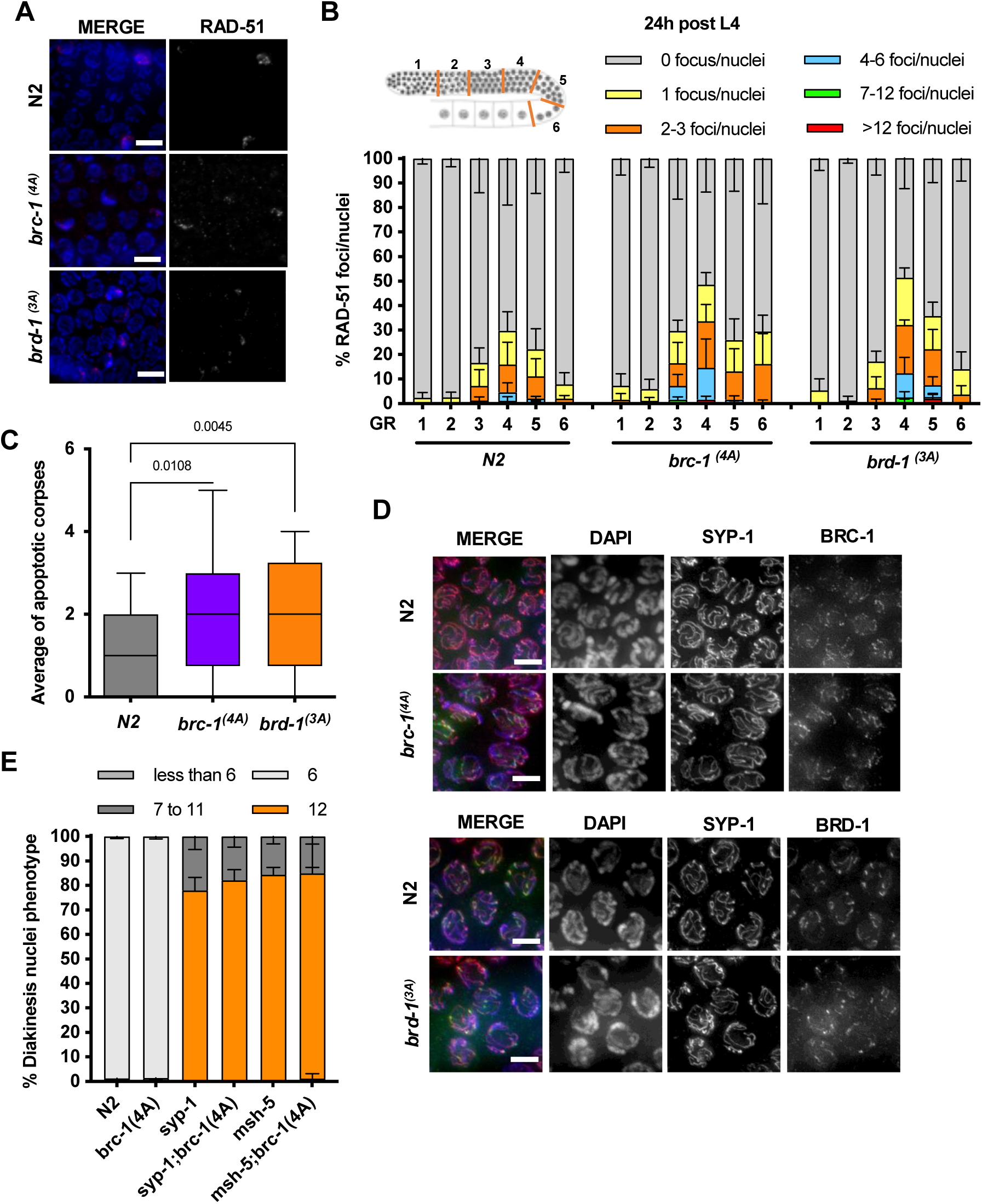
*brc-1* and *brd-1* phospho-mutants show mild defects on meiosis DSB repair. **(A)** Quantification of recombination marker RAD-51 foci in the indicated strains in normal conditions. A minimum of 4 gonads from 4 independent experiments were analyzed and 70-80 nuclei were scored for each region per gonad. On the top diagram of a hermaphrodite gonad, indicating the regions in which the number of RAD-51 foci was scored: 1) mitotic division zone; 2) transition zone; 3) early pachytene; 4) mid-pachytene; 5) late pachytene; 6) diplotene. Data are represented as mean ± SD **(B)** Representative images of mid-pachytene nuclei (region 4) stained with anti-RAD-51 (red) and DAPI (blue) for the indicated genotypes. Scale bar, 5µm. **(C)** Quantification of apoptotic corpses stained with SYTO-12 at late pachytene stage in the indicated strains in normal conditions. A minimum of 30 gonads from 4 independent experiments were analyzed. Data are represented as mean ± SD and p-value for unpaired t-test are indicated. **(D)** Representative images of mid-pachytene stage nuclei stained with the antibody against synaptonemal complex protein SYP-1 (red), BRC-1 (green on top) or BRD-1 (green on bottom) and counterstained with DAPI for the indicated genotypes. Scale bar, 3µm. **(E)** Quantification of the number of DAPI-stained bodies in the diakinetic oocytes of the indicated strains in normal conditions. -4 to -1 oocytes from a minimum of 10 gonads from 3 independent experiments were analyzed for each strain and condition. Data are represented as mean of the different experiments ± SEM.

Finally, it has been shown that both proteins BRC-1 and BRD-1 are dependent on each other for mutual stability and co-localize on the SC during prophase I in *C. elegans* germline (11). Since BRC-1/BRD-1 localization is essential for DSB repair upon induction of exogenous damage during gametogenesis, we investigated the localization of both proteins in our mutant alleles. Detection was performed by employing previously characterized antibodies against BRC-1 and BRD-1 (40). Immunofluorescences with αBRC-1 in *brc*-*1*^4A^ germlines or αBRD-1 in *brd*-*1*^3A^ germlines showed that both proteins localize at meiotic chromosomes with a pattern similar to that of the SC (Figure 2D), as previously described (11). To further proof that our alanine mutants are not behaving as a null mutation we generated double mutans of *brc*-*1*^4A^ with *syp-1* and *msh-5*. It has been shown that loss of *brc-1* in backgrounds where CO formation is prevented leads to chromosome fragmentation at diakinesis (12), contrary our double mutants show the expected 12 univalents at diakinesis (Figure 2E), supporting that the phenotypes we are observing are due to the impossibility to phosphorylate BRC-1 and not to defects in the functionality of the protein.

#### Loss of BRC-1 and BRD-1 phosphorylation leads to IR sensitivity

Since we observed phosphorylation of BRC-1 and BRD-1 under DNA damage conditions, we next analysed whether the phosphorylation status of BRC-1 or BRD-1 impacts the ability of nematodes to respond to exogenous DNA damage induced by IR. First, we determined IR sensitivity by scoring survival of the resulting F1 progeny 24–36 h after irradiation of L4 stage hermaphrodites with different doses (31). Irradiation with low doses resulted in a slight decrease in survival of 10% in N2(wt) and *brd-1^3A^*allele and 17% in *brc-1^4A^* mutant allele (Figure 3A). However, at higher doses (75Gy and 100Gy), both mutant alleles exhibited reduced survival relative to the N2(wt) strain (Figure 3A). This data indicates that the strains containing the non-phosphorylatable alleles of *brc-1* and *brd-1* are more sensitive to IR.

**Figure 3.**
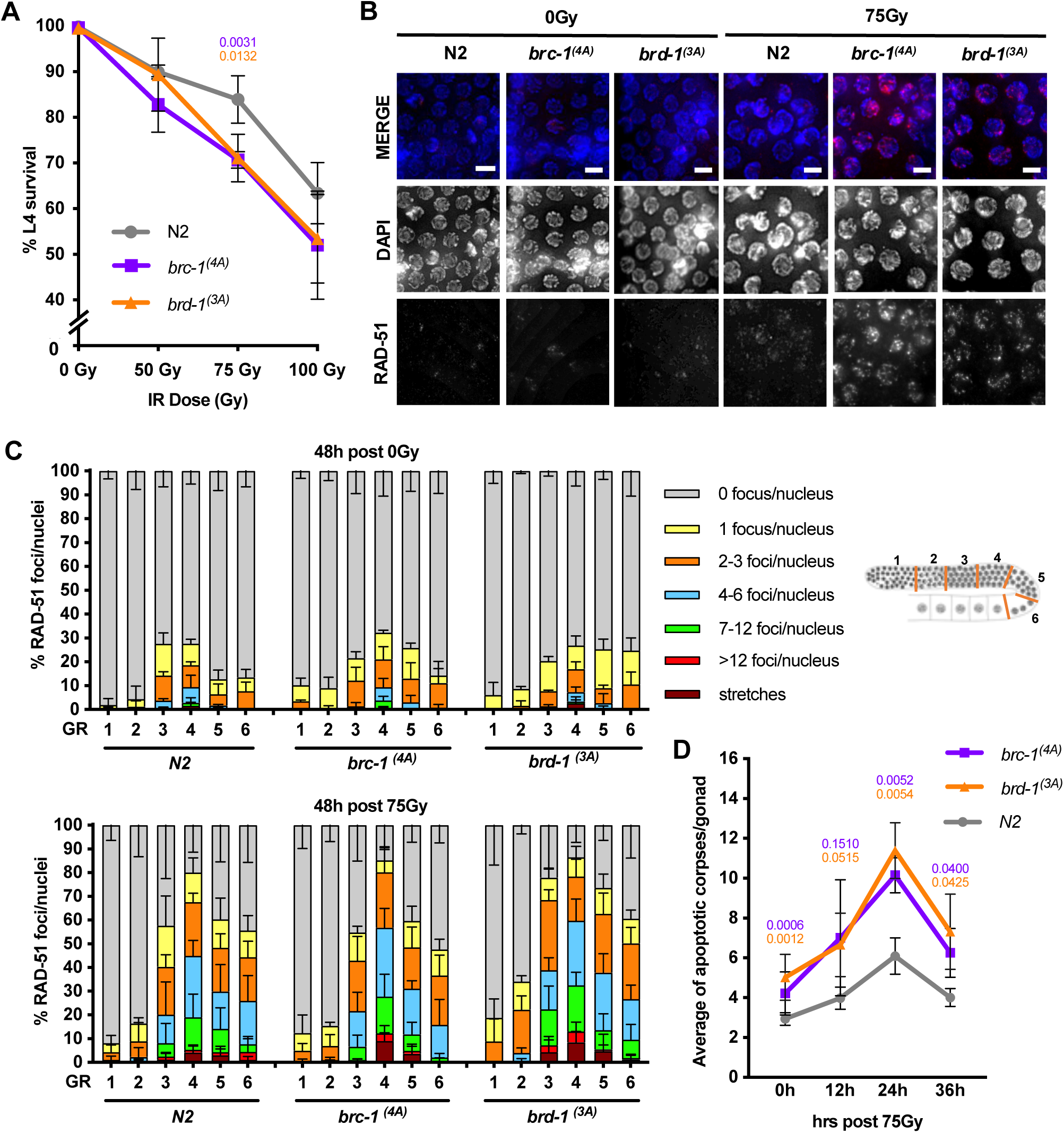
Loss of BRC-1 and BRD-1 phosphorylation leads to IR sensitivity. **(A)** Sensitivity of L4-stage worms from the indicated strains to different doses of irradiation (IR). Survival percentage of offspring is shown. Data are represented as average percentage ± SD from 3 independent experiments with 15 worms each. P-values for unpaired significative t-test are indicated. **(B)** Representative images of mid-pachytene nuclei stained with anti-RAD-51 (red) and DAPI (blue) for the indicated genotypes, 48 h after irradiation at doses indicated. Scale bar, 5µm. **(C)** Quantification of recombination marker RAD-51 foci in the indicated strains 48 h after irradiation at doses indicated on top of each graph. A minimum of 3 gonads from 3 independent experiments were analyzed and 70-80 nuclei were scored for each region per gonad. On the right, a diagram of a hermaphrodite gonad, indicating the regions in which the number of RAD-51 foci was scored: 1) mitotic division zone; 2) transition zone; 3) early pachytene; 4) mid-pachytene; 5) late pachytene; 6) diplotene. Data are represented as average ± SD. **(D)** Quantification of apoptotic corpses stained with SYTO-12 in late pachytene stage in the indicated strains at different times after 75Gy irradiation. A minimum of 20 gonads from 4 independent experiments were analyzed. Data are represented as average ± SD and p-value for significative unpaired t-test are indicated.

We showed above that *brc*-*1^4A^* and *brd*-*1^3A^*show mild defects in repairing programmed meiotic DSBs. To determine how *brc*-*1^4A^*and *brd*-*1^3A^* alleles respond to an excess of DSBs we analysed the intensity and distribution of RAD-51 foci 48h post-treatment with 75Gy IR since at 24h we still saw mitotic cell cycle arrest. Under non-treated conditions we could observe similar results as before, in which both mutants show modestly increased RAD-51 foci until late pachytene when compared with the N2(wt), suggesting a defect in DSB repair (Figure 3B, 3C, Supplementary Figure S2). In germlines from N2(wt) treated worms, we observed a mild increase in the levels of RAD-51 at the transition zone and an increase in the percentage of nuclei with RAD-51 foci compared to germlines from non-irradiated N2(wt) worms (Figure 3B, 3C, Supplementary Figure S2), implying that at this time point the N2(wt) has less DSB repair efficiency. Both non-phosphorylatable mutant strains also showed a substantial elevation in RAD-51 signal in all the germline regions after irradiation (Figure 3B, 3C, Supplementary Figure S2). In contrast with the N2(wt) we observe more nuclei in the category of <12 foci or RAD-51 stretches in the *brc*-*1*^4A^ and *brd*-*1*^3A^ mutant strains. We also quantified apoptotic corpses at different time points after L4 stage hermaphrodites IR treatment. As can be observed in the graph, *brc*-*1*^4A^ and *brd*-*1*^3A^ mutant strains showed a significant increase of apoptosis throughout the time course compared to the control N2(wt) (Figure 3D). This data strongly indicates that the repair of excess meiotic DSBs requires phosphorylation of BRC-1 and BRD-1 proteins.

#### IR-dependent meiosis catastrophe in non-phosphorylatable *brc-1* and *brd-1* alleles

While performing the RAD-51 analysis of L4 stage hermaphrodites at 48hr after irradiation, we noticed an increase in diakinesis nuclei with abnormalities. In *C. elegans* successful meiotic recombination and crossover formation results in 6 bivalents at diakinesis however mutants defective in DSB repair usually show altered DAPI bodies at diakinesis, particularly after exogenous DNA damage. Therefore, we analysed the DNA structures of diakinesis nuclei along the z-stacks of the acquired images (a 3D reconstruction is included in Figure 4A and videos of several examples can be found in supplemental material). In unchallenged conditions, DAPI staining of diakinesis cells showed no significant differences between the N2(wt) and *brc*-*1*^4A^ and *brd*-*1*^3A^ mutant strains: the majority of diakinesis presented 6 DAPI stained bodies (Figure 4A, B). In a few cases we observed chromosome fusions in nuclei of the phospho-mutant alleles (Figure 4B). In all the strains, including the N2(wt), we detected an increase in aneuploid diakineses following IR treatment (75Gy) (Figure 4A, B). Strikingly, however, *brc*-*1*^4A^ and *brd*-*1*^3A^ mutants’ strains showed a dramatic increase in chromosome aberrations, including altered numbers of DAPI bodies (Figure 4B), chromosome fusions, fragmentation or both (Figure 4A panels a-c-d, 4B). In addition, 10.8% of *brc*-*1*^4A^ and 5.8% of *brd*-*1*^3A^ diakinesis nuclei exhibit diffuse chromosome masses (Figure 4A panel b, 4B). This observation indicates that failure to repair excess meiotic DSBs results in aneuploidy when BRC-1 and BRD-1 cannot be phosphorylated.

**Figure 4.**
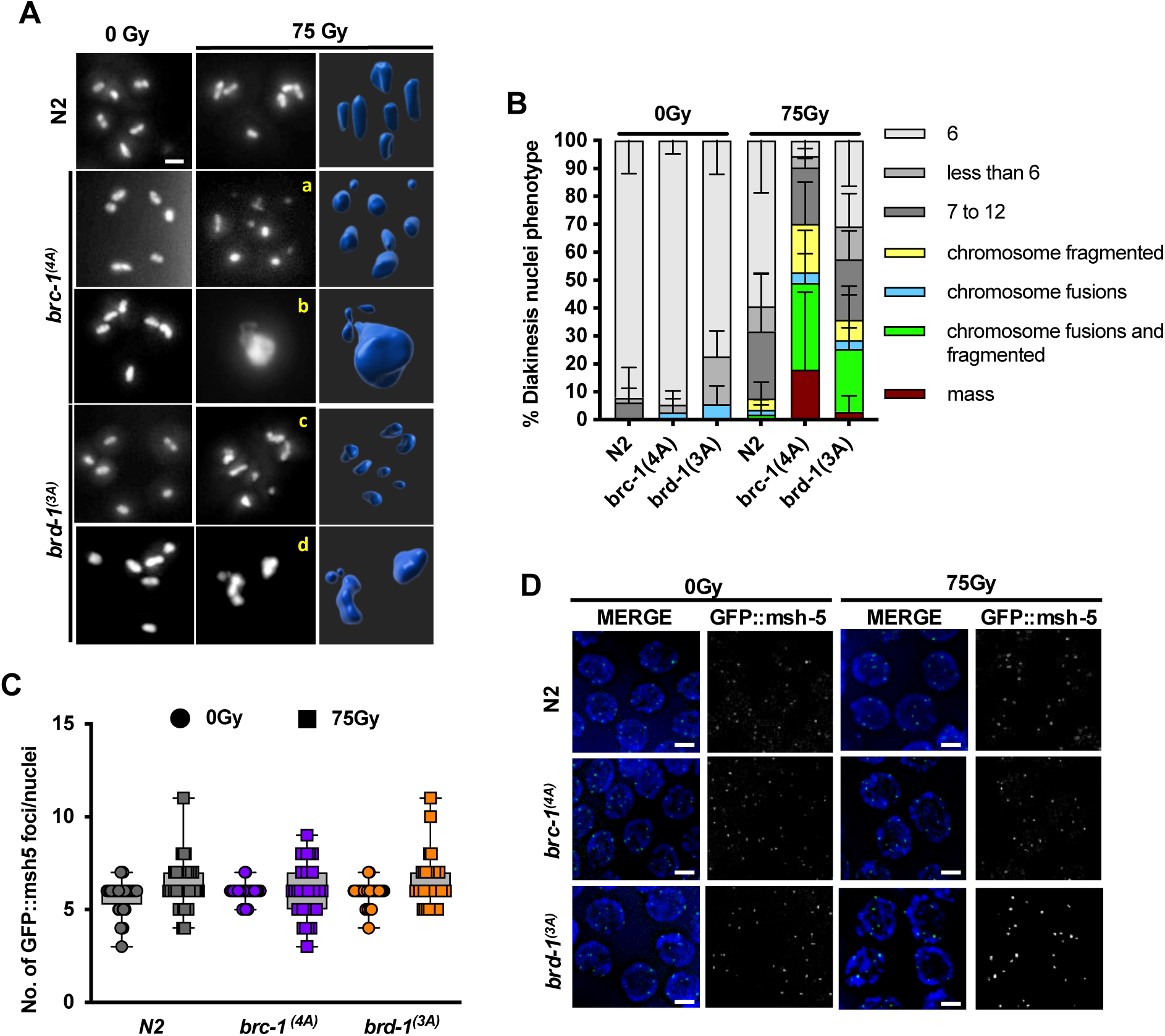
IR-dependent meiosis catastrophe in non-phosphorylatable *brc-1* and *brd-1* alleles. **(A)** Representative diakinesis-stage nuclei of the indicated strains and conditions stained with DAPI showing normal karyotype (6 bivalents) and defective diakinesis products: a) chromosome fragmentation; b) mass; c) chromosome fusions and fragmentation; d) chromosome fusions 48 h after 75Gy irradiation. Scale bar, 3 µm. At right, Imaris Surfaces model for object detection from the correspondent diakinesis nuclei showed. **(B)** Quantification of the different diakinesis nuclei phenotype in the diakinetic oocytes of the indicated strains in normal conditions and 48 h after 75 Gy irradiation. -4 to -1 oocytes from a minimum of 10 gonads from 3 independent experiments were analyzed for each strain and condition. Data are represented as the mean of the different experiments ± SEM. **(C)** Quantification of pro-crossover marker GFP::MSH-5 foci in the indicated strains 48 h after 75Gy irradiation. A minimum of four gonads from four independent experiments were analyzed and 12 nuclei from late pachytene stage were scored per gonad. Individual values are represented. P-values for unpaired t-test weren’t significative. **(D)** Representative images of late-pachytene stage nuclei stained with pro-crossover marker MSH-5 fussed to GFP (green) and DAPI (blue) for the indicated strains 48 h after 75Gy irradiation. Scale bar, 5 µm.

The high percentage of diakinesis nuclei with abnormal chromosomes did not correlate with the survival curves we observed in the IR sensitivity assay. Is important to notice that the experiments are not performed at the same developmental stage and time after IR treatment. The sensitivity assay analyses the 24h post irradiation progeny of L4 stage hermaphrodites while the IFs are done 48hr post irradiation of 24h post L4 stage hermaphrodites. Therefore, we carry out an IR sensitivity assay in which we analysed the 48h post irradiation progeny of L4 stage hermaphrodites. Surprisingly the survival of the N2(wt) in this condition is severely affected as well as in both phospho-alleles strains (Supplementary Figure S3A). We noticed that in the post 24hr assay the brood size at after IR treatment halves while in the post 48hr is not the case (Supplementary Table S1). Thus, this could be in part related with the age of the nematodes and could be masking the data.

#### Phosphorylation of BRC-1 regulates HR intermediate processing

Repair of meiotic DSB is crucial for proper formation of crossovers (CO); *C. elegans* exhibit extreme interference with only one CO formed per homolog pair. It has been shown that loss of *brc-1* and *brd-1* does not affect the number of CO but leads to an altered CO landscape (10). Nevertheless we checked the number of CO in the phospho-mutant alleles, to this end we combined the alleles with the CO marker *gfp::msh-5* (11) and analysed MSH-5 foci, a marker of COs. In normal conditions, we observed six CO in N2(wt), *brc*-*1*^4A^ and *brd*-*1*^3A^ strains at late pachytene. In germlines challenged with IR we observed no significant increase in the number of MSH-5 foci in N2(wt) worms and the same was observed in the strains carrying the non-phosphorylatable alleles (Figure 4C, D). This data indicates that the number of CO is not overtly affected in *brc*-*1*^4A^ and *brd*-*1*^3A^ even if excess DSBs are introduced by IR and inter-homolog recombination is preserved in our mutants’ alleles.

In *C. elegans* the NHEJ repair pathway requires the activities of the CeKU70-CeKU80 heterodimer and *lig-*4 (Ligase IV). Previous studies have shown that illegitimate activation of NHEJ is in part responsible for the meiotic abnormalities observed in HR mutants (41). Having observed that IR-dependent DSBs lead to defective diakinesis in the phospho-mutant strains, we wanted to address whether NHEJ contributes to this phenotype. To this end, we analysed diakinesis nuclei with and without IR in N2(wt), *brc*-*1*^4A^ and *brd*-*1*^3A^ strains subject to *lig-4* RNAi. Under normal conditions, N2(wt), *brc*-*1*^4A^ and *brd*-*1*^3A^ strains worms presented with six bivalents at diakinesis, and this was modestly altered by *lig-4 RNAi*. After IR treatment, the presence of aberrant chromosomal fusion and fragmentations in the *brc*-*1*^4A^ and *brd*-*1*^3A^ germlines were exacerbated irrespective of the status of LIG-4 (Figure 4A). These data indicate that NHEJ does not make a significant contribution to the improper repair of IR-dependent DSBs in these strains, suggesting that alternative DSB repair processes are driving mis-repair in the absence of phosphorylation of BRC-1 and BRD-1.

It has been proposed that TMEJ acts as an alternative repair mechanism to HR in the *brc- 1* mutant (14, 16). To test the involvement of TMEJ, we generated double mutants of *brc*-*1*^4A^ and *polq-1*. While we observed no overt differences in the *polq-1*;*brc*-*1*^4A^ strain between the single mutants in the absence of IR (Figure 5B, 5C), IR treatment lead to a synergetic effect in the *polq-1*;*brc*-*1*^4A^ strain. Analysis of the double mutant *polq-1*;*brc*-*1*^4A^ strain showed that IR treatment led to a general increase of fragmentations and fusions compared with the single mutants. (Figure 5B, 5C). These results indicate that TMEJ is not responsible for the abnormal repair of IR-dependent DSBs in the *brc*-*1*^4A^ strain.

**Figure 5.**
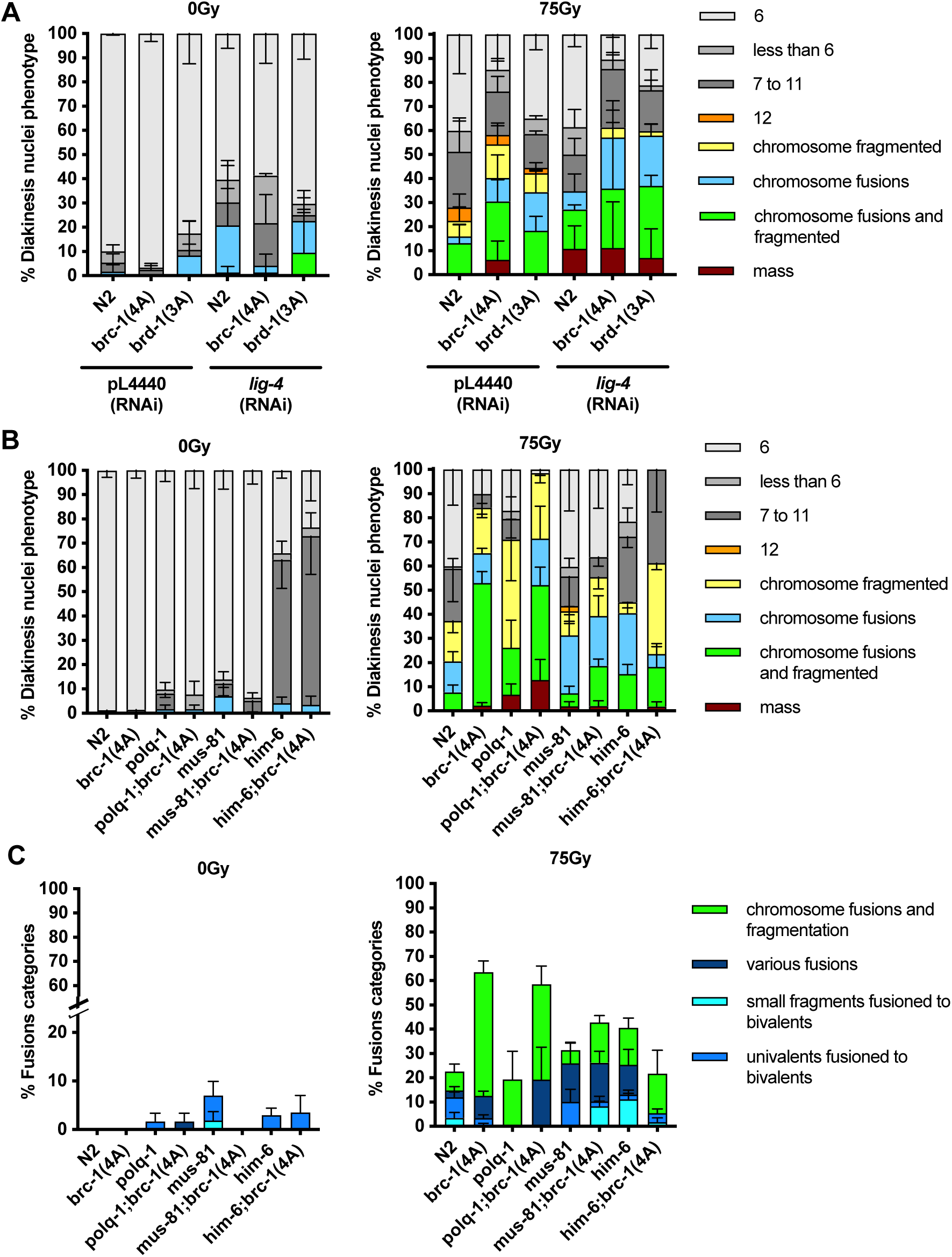
Meiosis catastrophe of the phospho-mutants is dependent on HIM6/MUS81. **(A)** Quantification of the different diakinesis nuclei phenotype in the diakinetic oocytes of the indicated strains with/without LIG-4 RNAi depletion in normal conditions and 48 h after 75Gy irradiation. -4 to -1 oocytes from a minimum of 10 gonads from 3 independent experiments were analyzed for each strain and condition. Data are represented as the mean of the different experiments ± SEM. **(B)** Quantification of the different diakinesis nuclei phenotype in the diakinetic oocytes of the indicated strains 48 h after the irradiation dose indicated on top of each graph. -4 to -1 oocytes from a minimum of 10 gonads from 3 independent experiments were analyzed for each strain and condition. Data are represented as mean of the different experiments ± SEM. **(C)** Percentage of diakinetic oocytes of the indicated strains showing fusions in normal conditions (left) and 48 h after 75Gy irradiation (right). -4 to -1 oocytes from a minimum of 10 gonads from 3 independent experiments were analyzed for each strain and condition. Data are represented as mean ± SEM.

Although COs are not affected in our phospho-alleles, it has been shown that BRC-1 is required for efficient processing of recombination intermediates. We therefore examined diakinesis in the absence of later HR intermediate processing factors, including the resolvase MUS-81 and Holliday junction resolution factor HIM-6 (Bloom). Again, no changes were observed in the double mutants in the absence of DNA damage treatment. In contrast, in gonads of *mus- 81*;*brc*-*1*^4A^ and *him-6*;*brc*-*1*^4A^ exposed to IR we observed a suppression in chromatin masses, more robustly in the *him-6* double mutant, as well as a reduction in chromosome fusions and an increase in fragmentation (Figure 5B, 5C). Moreover, in the double mutant with *him-6* the reduction in aberrations results in an increase in the univalents after IR (Figure 5B). This data reveal that the phosphorylation of BRC-1/BRD-1 after IR is important for the correct regulation of HR intermediate resolution/dissolution.

The reduction in diakinesis fusions when combining *brc*-*1*^4A^ allele with the resolvases prompt us to check if this could alleviate the IR sensitivity observed in the non-phosphorylatable *brc-1* allele. For that we included the *mus-81*;*brc*-*1*^4A^ strain in the sensitivity assay performed at 48h post irradiation progeny of L4 stage hermaphrodites. First we found that *mus-81* is extremely sensitive to IR (Supplemental figure S3A), additionally the brood size reduction was exacerbated and even more in the *mus-81*;*brc*-*1*^4A^ mutant (Supplemental figure S3B). However in the *mus- 81*;*brc*-*1*^4A^ strain we detected an increase of survival to *the single mutant brc*-*1*^4A^ levels. This data suggests that both act in the same pathway to deal with excessive DNA damage.

## Discussion

In response to exogenous DSBs, the cellular response to DNA damage is initiated by the activation of ATM/Tel1 and ATR/Mec1 kinases, which are conserved throughout eukaryotes (42, 43). In *C. elegans* it has been shown that both kinases also regulate key steps of meiosis, phosphorylating several target proteins involved in DSB formation and repair, cohesion and synapsis between chromosomes (23–26, 44). In previous work from our group, we described that IR-induced phosphorylation of SYP-1, a central synaptonemal complex component, is required for the repair of excessive meiotic DSBs relying on *brc-1* activity (25), and it has been recently shown that BRC- 1 regulates homolog-independent processing (13). In this work we describe damage-induced phosphorylation of BRC-1/BRD-1, which we propose impacts template bias by controlling HR intermediates resolution. In contrast to *brc-1/brd-1* null mutants, the non-phosphorylatable BRC-1 and BRD-1 alleles do not result in an increase of males in the progeny nor overt defects in meiotic DSB repair in the scenario where CO is impeded, indicating that DSB repair through the homolog is not compromised in the phospho-alleles (11, 12, 35). However, *brc*-*1*^4A^ and *brd*-*1*^3A^ strains do exhibit a robust DNA damage sensitivity phenotype, which we attribute to DSBs repair defects that lead to meiotic catastrophe and aneuploidy at diakinesis.

Similar to their human counterparts, *C. elegans* BRC-1 and BRD-1 proteins interact through the N-terminal RING finger domains (40, 45). Notably, the phosphorylation site identified in the BRD-1 protein, a cluster of three TQ, is located adjacent to the RING domain. Despite the proximity to the RING domain, phosphorylation of the TQ cluster is unlikely to affect heterodimerization as the mutant strains exhibited normal BRC-1/BRD-1 localization on the SC. Indeed, it has been described that the BRC-1/BRD-1 heterodimer is interdependent in their localization along the SC and that the loss of either protein in synapsis deficient backgrounds leads to embryonic lethality (10). The phospho-site of BRC-1 is located in one of the two BRCT repeat domains. BRCT domains are found in many DNA replication and repair proteins, are required to bind phosphorylated proteins, and are also involved in DNA and PAR binding (46). In the case of human BRCA1, the BRCT domain is essential for tumour suppressor activity (47). While the precise function of phosphorylation of BRC-1 within the BRCT domain is unknown, it is tempting to speculate that this may modulate, positively or negatively, BRC-1’s ability to bind a substrate important for repair.

Previous research has suggested that BRC-1 activity can stimulate non-crossover-mediated repair, and that in its absence, alternative repair mechanisms may be activated to repair intermediates (12, 48). BRCA1 has been shown to suppress NHEJ in order to promote the repair of DSBs through HR. Since NHEJ has been linked to meiotic aberrations observed in HR mutants, we hypothesized that NHEJ repair may be used as a last resource to deal with the additional damage caused by IR in our phospho-mutants. However, contrary to our expectations, aberrant meiotic diakinesis were exacerbated after IR in the phospho-mutants lacking *lig-4*. We believe that while the inactivation of NHEJ is dependent on BRCA1, this role does not rely on these BRC-1 and BRD-1 specific phosphorylation sites. We also examined alternative non-homologous end joining, also known as TMEJ that functions to repair resected DSBs in the absence of HR (49), and it has been shown to be the source of genome structural variations in a BRCA1 or BARD1 deficient scenario (14). Similarly to what we observed with NHEJ, TMEJ loss in the phospho-mutants conferred an increase in chromosomal abnormalities in IR treated animals. These data indicate that neither NHEJ nor TMEJ are responsible for the abnormal diakinesis we observe.

Meiotic DSBs repair via HR is believed to results in the formation of double HJ (dHJ) intermediates, which can be resolved through cleavage by HJ resolvases, which results in either a crossover (CO) or non-crossover (NCO) product or dissolution by the BTR complex, consisting of the HIM-6/BLM helicase, topoisomerase IIIα, RMI1, and RMI2 (50, 51). Importantly, CO designation is unaltered in the phosphomutants when there is excess DNA damage. In *C. elegans* it has been proposed that MUS-81 and HIM-6 act redundantly to resolve joint molecules early in meiosis, presumably to form NCO (5, 6, 52). Strikingly, we show that the aberrant meiotic chromosomes and fusion observed in the BRC-1 phospho-mutants after IR treatment, are largely suppressed by removing HIM-6 and MUS-81.

In summary, we have identified DNA damage dependent phosphorylation events in BRC- 1 and BRD-1, which protect germline cells from meiotic catastrophe upon IR treatment. We have excluded that this phosphorylation is to control alternative repair pathways as NHEJ or TMEJ, since the double mutans do not suppress the abnormal chromosomal rearrangements that we observe in the non-phosphorylatable mutants. Unexpectedly we have uncovered a role of BRC-1 in balancing the resolution/dissolution of recombination intermediates between homolog chromosomes, since removal of HIM-6/BLM or MUS-81 suppresses the aberrant meiotic chromosomes and fusions of the *brc-1^4A^* mutant. Hence, we propose that phosphorylation of BRC- 1/BRD-1 by ATM and ATR in response to excess DNA damage regulates HR intermediate processing by MUS-81 and HIM-6/BLM. Our data support a model (Figure 6) in which damage-induced BRC-1/BRD-1 phosphorylation is involved in the processing of NCOs. In this way, the cell regulates the excess inter-homolog crossovers, which could hinder correct chromosome segregation at the first meiotic division. Considering the conserved nature of most repair processes across species, it is possible that BRCA1 and BARD1 play similar roles in meiosis and DNA repair in higher eukaryotes. Indeed, the protein sequence of the human orthologs contain similar S/T-Q motifs. It will also be interesting to consider if this regulatory mechanism is sex-specific since it has been shown in the nematode that BRC-1/BRD-1 differently regulates meiotic DSB repair and CO determination in male and female meiosis (16).

**Figure 6.**
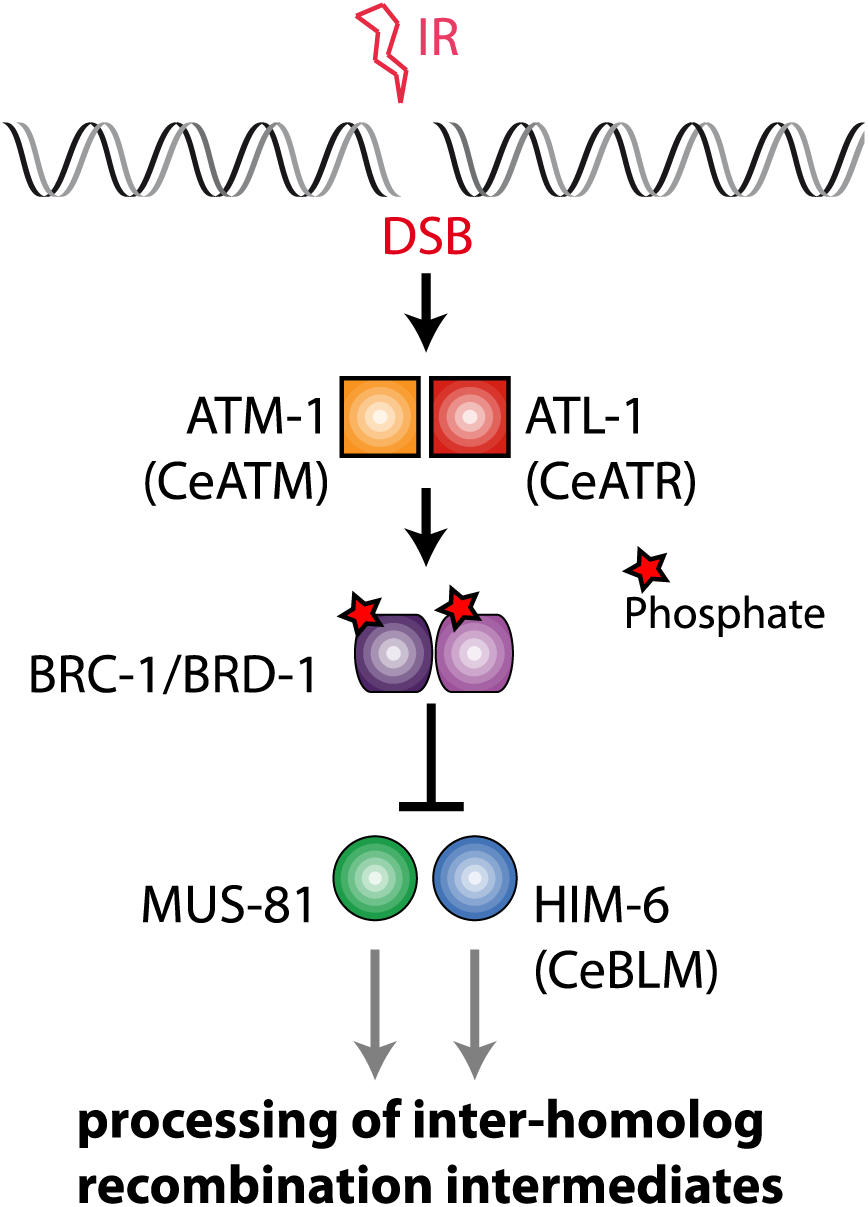
Proposed model. BRC-1/BRD-1 phosphorylation dependent of ATM-1/ATL-1 regulate HIM-6/MUS-81 function, correctly balancing the resolution/dissolution of recombination intermediates arising between homolog chromosomes. Through this mechanism, the cell regulates the processing and repair of excess inter-homolog crossovers, which could hinder correct chromosome segregation at the first meiotic division.

**Supplementary Figure S1. SYP-1 assembly in *brc-1^(4A)^*and *brd-1^(3A)^ C. elegans* mutants.** Representative images of fixed germlines of indicated genotype in normal conditions immunostained with anti-SYP-1 antibody and counterstained with DAPI. Scale bar 15μm. At right, zoom-in of pachytene stage from the correspondent germline. Scale bar, 3µm. On top, scheme depicting the gonad regions: 1) mitotic division zone; 2) transition zone; 3) early pachytene; 4) mid pachytene; 5) late pachytene; 6) diplotene.

**Supplementary Figure S2. RAD-51 accumulation in *brc-1^(4A)^*and *brd-1^(3A)^* 48h after 75Gy irradiation.** Representative images of fixed germlines of indicated genotype in normal conditions and 48 h post 75Gy, immunostained with anti RAD-51antibody and counterstained with DAPI. Scheme depicting the gonad regions: 1) mitotic division zone; 2) transition zone; 3) early pachytene; 4) mid pachytene; 5) late pachytene; 6) diplotene. Scale bar 15μm.

**Supplementary Figure S3. IR-dependent analysis in non-phosphorylatable *brc-1* and *brd-1* alleles. (A)** Sensitivity of L4-stage worms from the indicated strains after 48h irradiation. Survival percentage of offspring is shown. Data are represented as average percentage ± SD from 3 independent experiments with 15 worms each. **(B)** Brood size of L4-stage worms from the indicated strains after 48h irradiation. Data is shown as average ± SD from independent experiments with 15 worms each. Also is shown the average of each experiment. P-values for Tukeys multiple comparison-test are indicated.

## Acknowledgments

The publication of this article is funded by Spanish Ministry of Science and Innovation, the Spanish Research Agency, and the European Regional Development Fund - Proyecto PID2021-123850N-I00 financiado por MCIN/ AEI /10.13039/501100011033/ y por FEDER Una manera de hacer Europa.

We thank S. Boulton and P. Huertas for comments in the manuscript, and V. Jantsch, M. Olmedo, P. Askjaer, P. Huertas for sharing antibodies and strains.

