## Supplemental for "Damage-induced phosphorylation of BRC-1/BRD-1 in meiosis preserves germline integrity"

**Fig. S1**

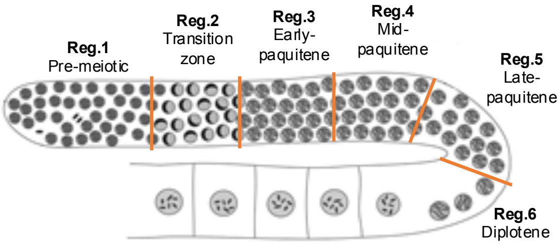

Reg.1
Pre-meiotic
Reg.2
Transition zone
Reg.3
Early-paquitene
Reg.4
Mid-paquitene
Reg.5
Late-paquitene
Reg.6
Diplotene


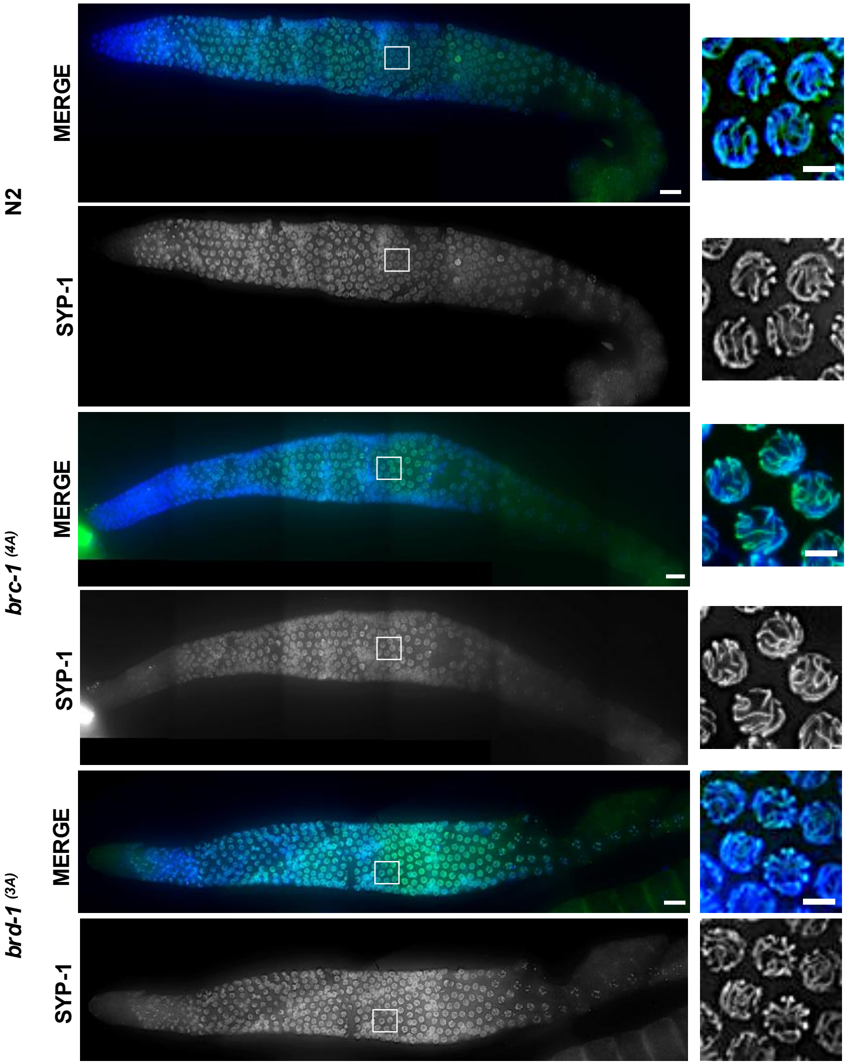

N2
MERGE
SYP-1
brc-1 (4A)
MERGE
SYP-1
brd-1 (3A)
MERGE
SYP-1

**Fig.S2**

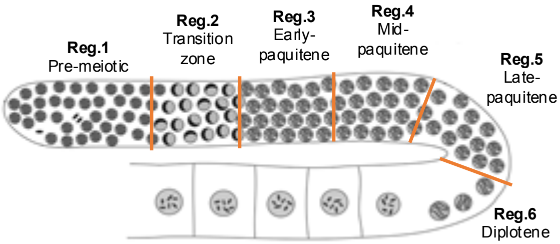
Reg.1
Pre-meiotic
Reg.2
Transition zone
Reg.3
Early-paquitene
Reg.4
Mid-paquitene
Reg.5
Late-paquitene
Reg.6
Diplotene

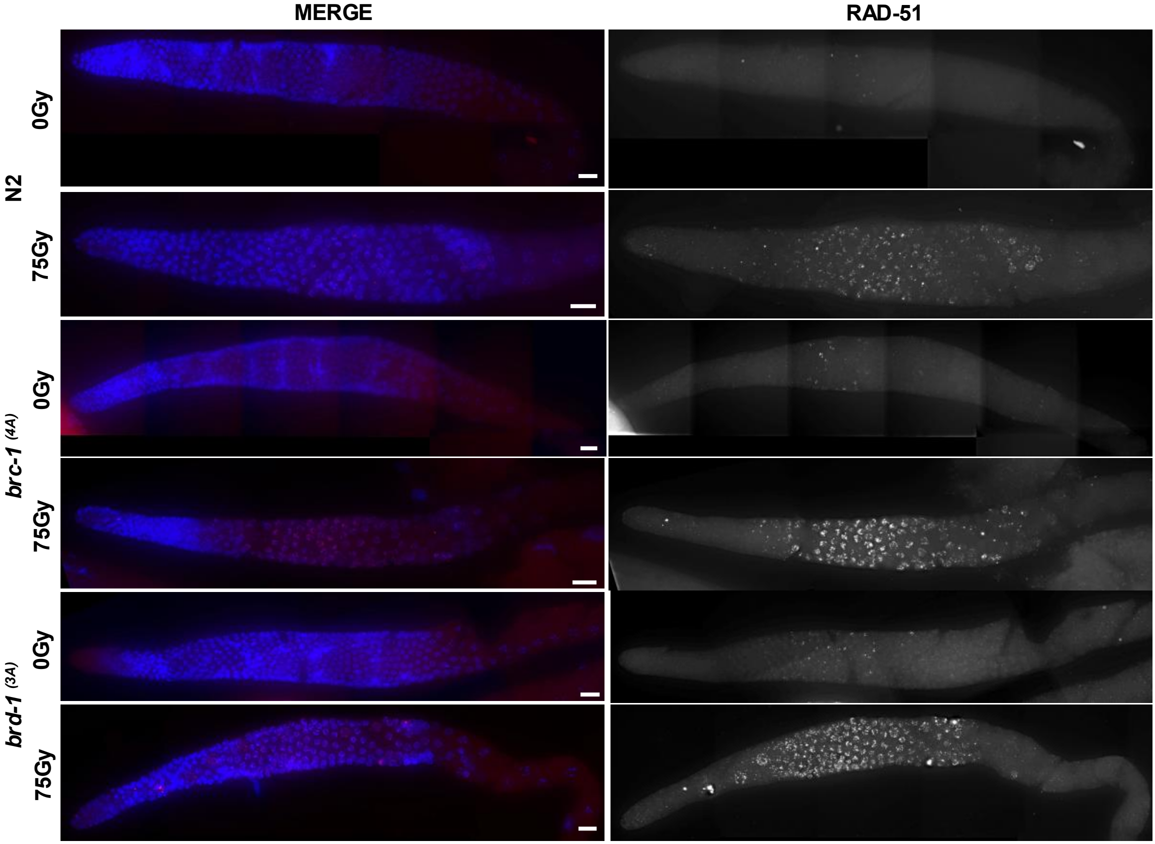
MERGE
RAD-51
N2
0Gy
75Gy
brc-1 (4A)
0Gy
75Gy
brd-1 (3A)
0Gy
75Gy

**Fig. S3**

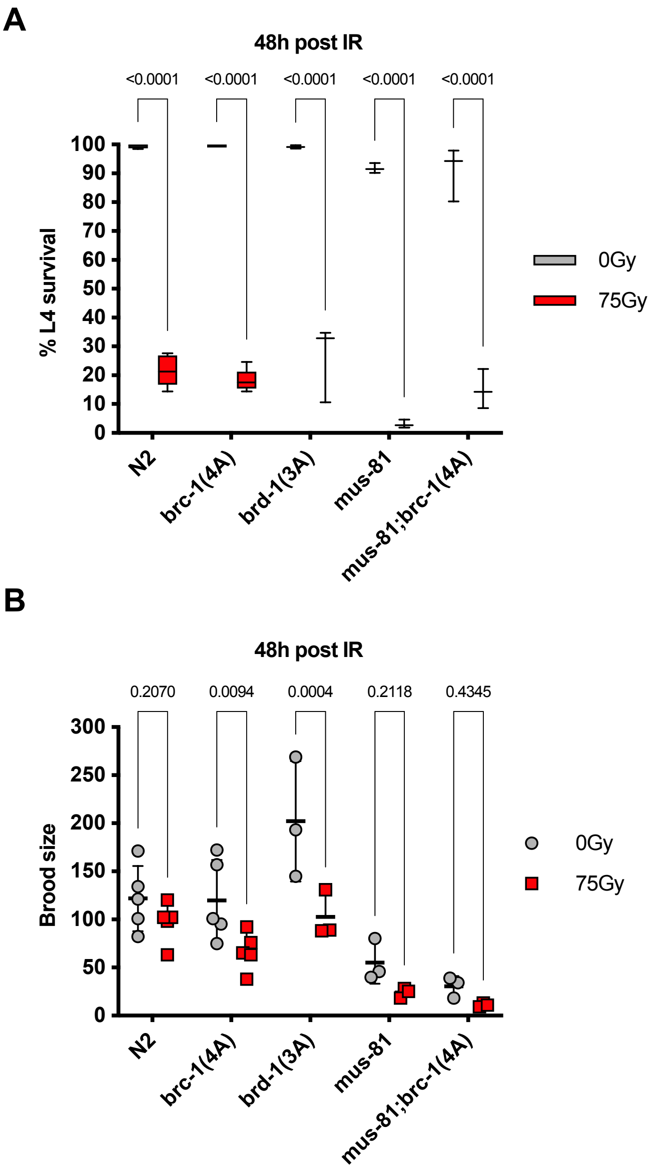
A
48h post IR
<0.0001
<0.0001
<0.0001
<0.0001
<0.0001
% L4 survival
100
90
80
70
60
50
40
30
20
10
0
N2
brc-1(4A)
brd-1(3A)
mus-81
mus-81;brc-1(4A)
0Gy
75Gy
B
48h post IR
0.2070
0.0094
0.0004
0.2118
0.4345
Brood size
300
250
200
150
100
50
0
N2
brc-1(4A)
brd-1(3A)
mus-81
mus-81;brc-1(4A)
0Gy
75Gy

| **Table S1. Brood size analysis of *brc-1*$^{(4A)}$ and *brd-1*$^{(3A)}$ mutant alleles** | | | | |
|---|---|---|---|---|
| | **Average Brood ± SD** | | | |
| | **24h post L4 and IR** | | **48h post L4 and IR** | |
| **Strain** | **0Gy** | **75Gy** | **0Gy** | **75Gy** |
| N2 (WT) | 337.17 ± 36.07 | 114.08 ± 27.12 | 141.40 ± 45.04 | 104.60 ± 7.85 |
| *brc-1*$^{(4A)}$ | 301.08 ± 58.74 | 111.33 ± 21.88 | 159.8 ± 36.37 | 78.2 ± 27.09 |
| *brd-1*$^{(3A)}$ | 287.33 ± 38.51 | 117.70 ± 20.27 | 120.89 ± 63.27 | 90.56 ± 35.12 |

| **Table S2. *C. elegans* strains** | | |
|---|---|---|
| **Strain** | **Genotype** | **Source** |
| N2 | Wild type Bristol | CGC |
| PHX4192: *brc-1*$^{(4A)}$ | brc-1(syb4192) III | This study |
| GIN127: *brd-1*$^{(3A)}$ | brd-1(3A) III | This study |
| 1960 VJ | ollas::cosa-1 III;msh-5::GFP IV | V. Jantsch |
| GIN126 | brc-1(syb4192) III;msh-5::GFP IV | This study |
| GIN128 | brd-1(3A) III;msh-5::GFP IV | This study |
| 1147 VJ | him-6(ok412) IV/nT1 (IV;V) | V. Jantsch |
| 1163 VJ | mus-81(tm1937) I | V. Jantsh |
| 1533 VJ | polq-1(tm2026) III | V. Jantsh |
| AV307 | syp-1(me17) V/nT1[unc-?(n754) let-? qIs50] (IV;V) | A.Villeneuve |
| GIN138 | msh-5(me23)IV/nT1[unc-?(n754)let-? qls50] (IV;V) | This study |
| GIN129 | brc-1(syb4192) (III);syp-1(me17) V/nT1[unc-?(n754) let-? qIs50] (IV;V) | This study |
| GIN131 | mus-81(tm1937) I;brc-1(syb4192) III | This study |
| GIN133 | brc-1(syb4192) (III);him-6(ok412) IV/nT1 (IV;V) | This study |
| GIN136 | brc-1(syb4192) III;polq-1(tm2026) III | This study |
| GIN139 | brc-1(syb4192) (III);msh-5(me23)IV/nT1[unc-?(n754)let-? qls50] (IV;V) | This study |

| **Table S3. Reagents and resources** | | |
|---|---|---|
| **Reagent or resource** | **Source** | **Identifier** |
| **Antibodies** | | |
| Rabbit RAD-51 Antibody | Novus Biologicals | NB100-148 |
| Guinea pig SYP-1 Antibody | The A. Villenueve Lab | MacQueen et al., 2002 |
| Alexa Fluor Goat anti-Guinea Pig::488 | Molecular probes | A11073 |
| Alexa Fluor Goat anti-Rabbit::568 | Molecular probes | A11011 |
| **Chemicals, Peptides and Recombinant Proteins** | | |
| Vectashield | Vector laboratories | H-1000 |
| 4,6-Diamino-2-phenylindole dihydrochloride (DAPI) | Invitrogen | D1306 |
| GoTaq G2 Flexi DNA Polymerase | Promega | M7805 |
| PstI | Biolabs | R01405 |
| PvuII | Biolabs | R31515 |
| Hyp188I | Biolabs | R0617L |
| BstAPI | Biolabs | R0654S |
| **CRISPR-Cas9 reagents** | | |

| | | |
|---|---|---|
| Alt-R® S.p. Cas9 Nuclease | IDT | 1081058 |
| Alt-R® CRISPR-Cas9 tracrRNA | IDT | |
| Nuclease Free Duplex Buffer | IDT | |
| Alt-R® CRISPR-Cas9 *dpy-10* crRNA | IDT | |
| Ultramer® DNA Oligo *dpy-10(cn64)* | IDT | |
| **Protein 3D structures** | | |
| BRC-1 | AlphaFold | B6VQ60 |
| BRD-1 | AlphaFold | Q21209 |
| BRCA1/BARD1 RING domain | Brzovic et al., 2001 | P38398 - 1jm7 |
| BRCA1 BRCT domain | Williams et al., 2001 | P38398 - 1jnx |
| **Software and Algorithms** | | |
| CRISPOR | http://crispor.tefor.net/ | Concordet, J.P. et al., 2018 |
| Image J (FIJI) | https://imagej.net/Welcome | Schindelin et al., 2012 |
| Imaris 9.2 | https://imaris.oxinst.com/ | |
| Prism 9 | https://www.graphpad.com/scientific-software/prism/ | |
| AlphaFold | https://alphafold.ebi.ac.uk/ | Jumper et al., 2021<br>Varadi et al., 2021 |
| ChimeraX | https://www.cgl.ucsf.edu/chimerax/ | Pettersen et al.; 2021 |
| Leica Application Suite Advanced Fluorescence (LAS-AF) | Leica | N/A |
| Nikon Instruments Software (NIS) | Nikon | N/A |
| Zen 2 blue | Zeiss | N/A |
| **Other** | | |
| MEGAquick-spin$^{TM}$ plus Fragment DNA Purification Kit | iNtRON Biotechnology | 17290 |

| Table S4. Primers and sRNA designed for CRISPR | | |
|---|---|---|
| **Primer/sRNA** | **Sequence (5’-3’)** | **Use** |
| brd-1_seqE1 | CGCCACATTTCAACAGAAACCG | Genotyping and sequencing |
| brd-1_seqE2 | CGTGGAATATATCCAATTCTGGAG | Genotyping and sequencing |
| brc-1_seqE1 | GTTTTTTTCCCAGATATATTCGAAAAACCG | Genotyping and sequencing |
| brc-1_seqE2 | CGAATTATGCACAATCTCAAAAAACACCC | Genotyping and sequencing |
| brc-1 11Fw | GACCGTACACCGAAAGCAATTC | Genotyping |
| brc-1_seq-wt | ATAGAGATATGCTGTGGA | Genotyping |
| brc-1_seq-mutA | ATAGAGATATGCTGCTGC | Genotyping |
| BRC-1 E1 | CTCCGTAGCTTGAAGTCTCA | Genotyping |
| BRC-1 E2 | TTTCGGTGGCGCCACATGGA | Genotyping |
| BRC-1 I1 | AATATAGGCACCGGCGGGGA | Genotyping |
| BRC-1 I2 | TGTCGCATCGTCGGCATTAA | Genotyping |
| BRC-1 I3 | CTGACTGAAAATCATAGCGG | Genotyping |
| msh-5_E3 | CCCGATGAACCGAAATCGGTATGCG | Genotyping |
| msh-5_I4 | CTCTGATGTTCCTTCTTCTG | Genotyping |
| cosa-1_E3 | CGCACAGGTCAAAACTTTGTATTGG | Genotyping |
| cosa-1_I1 | CCATTTGTGCAATGCAATCCGTCC | Genotyping |
| polq-1_tm2026 E1 | AGTGTGCAAGTTTCGAAGAATCGC | Genotyping |
| polq-1_tm2026 E2 | GATGCTGAACGGACTAATTTAAGC | Genotyping |
| polq-1_tm2026 I1 | CGCCACGTTCATGTAGGATTCGGG | Genotyping |
| polq-1_tm2026 I2 | GTGAATTGCTAAAGGGCGTGCGCC | Genotyping |
| polq-1_tm2026 H2 | CAAGTACCGAATTCCTCGTG | Genotyping |
| mus-81_tm1937 E1 | CACGATCAATAGAGACTCTCCGCC | Genotyping |
| mus-81_tm1937 E2 | CTTTCTGCTCATCATATCTACCTCC | Genotyping |
| mus-81_tm1937 I1 | GTTAGGTATTTGGCAGACTTACCG | Genotyping |

| | | |
|---|---|---|
| mus-81_tm1937 I2 | TTGCGATGCTCACGATTGTCTGCG | Genotyping |
| mus-81_tm1937 H2 | GCAGAACATATTTGAGCTTCC | Genotyping |
| him-6_ok412 E1 | AATGGTCACGATGAAGAGCC | Genotyping |
| him-6_ok412 E2 | ACCGAATATAGCCGTTCGTG | Genotyping |
| him-6_ok412 I1 | ATCGACCATCAGAGAATCCG | Genotyping |
| him-6_ok412 I2 | TGCATCCTGCTCTGACATTC | Genotyping |
| msh-5 I3 | CGATATTTCAGAAGATCTTG | Genotyping |
| msh-5 I4 | CCTCTGATGTTCCTTCTTCTG | Genotyping |
| syp-1 SQ1 | CGGTGTTTGCGGCCTCCGCTC | Genotyping |
| syp-1 SQ2 | AGTTTTCCCTCTTCGAGCGC | Genotyping |
| brd-1crRNA-PAM216 | GCTCATATGCCAGACGAGAT (GGG) | crRNAguide sequence for 1st step |
| brd-1_RepT131A_F | CAACATCTCTAGCTCAAGCAGAGTTTGCCTGTGCACAAAACATTTTTAACAAAGAAAAAACTCCTGCAGATGCCGTAGAGAAATTCCTGGAGGCGCAAGCCCACATGCCAGATGAGTTCGCACAACTTGGTGAAGAGGATGATGATTTGATGTGTAAAGATGAAAACCGGTAAGTCGAAAAAATCTGCGTTCGGGGGGAA | Repair template 1st step |
| brd-1crRNA-PAM70 | AATTTCTCTACGGCATCTGC (AGG) | crRNAguide sequence for 2st step |
| brd-1_RepT104A_F | GTCGGAACAACATCTCTAgCTCAAGCAGAGTTTGCCTGTGCACAAAACATTTTTAACAAAGAAAAAACTCCaGCtGATGCCGTAGAGAAATTCCTGGAGGCGCAAGCCCACATGCCAGAT | Repair template 2st step |
| polq-1crRNA_PAM31rev | TATAGAATTTGTAATATGTA (CGG) | crRNAguide sequence |
| polq-1crRNA_PAM899fw | AAGGAATTGATGGACAGAGG (TGG) | crRNAguide sequence |
| polq-1_tgm2026 RT | CTTTCAAAAATTACTATTAAAACTTTTTCAAATTAAAAAATTTTAGAAATTCAAAAAAAAAATTTCTGAAAACATCACCGATATCTGGGGAGAGCTATATGGAATTGGTTGAAGTATCTGTTTTAAAAAATAAAGGAGCAGCTTTTAACATTTTTAAAACACTTG | Repair template |
